# Divalent metal ions boost effect of nucleic acids delivered by cell-penetrating peptides

**DOI:** 10.1101/2021.12.09.471911

**Authors:** Maria Maloverjan, Kärt Padari, Aare Abroi, Ana Rebane, Margus Pooga

## Abstract

Cell-penetrating peptides (CPPs) are promising tools for transfection of various substances, including nucleic acids, into cells. The aim of current work was to search for novel safe and effective approaches for enhancing transfection efficiency of nanoparticles formed of CPP and splice-correcting oligonucleotide (SCO) without increasing the concentration of peptide. We analyzed an effect of inclusion of calcium and magnesium ions into nanoparticles on CPP-mediated transfection in cell culture. We also studied the mechanism of such transfection as well as its efficiency, applicability in case of different cell lines, nucleic acid types and peptides, and possible limitations. We discovered a strong positive effect of these ions on transfection efficiency of SCO, that translated to enhanced synthesis of functional reporter protein. We observed significant changes in intracellular distribution and trafficking of nanoparticles formed with addition of the ions, without increasing cytotoxicity. We propose a novel strategy of preparing CPP-oligonucleotide nanoparticles with enhanced efficiency and, thus, higher therapeutic potential. Our discovery may be translated to primary cell cultures and, possibly, *in vivo* studies, in the aim to increase CPP-mediated transfection efficiency and likelihood of using CPPs in clinics.

## 1. Introduction

Gene therapy is a strategy of treatment or alleviating symptoms of disease by altering gene expression [1, 2] and is a rapidly growing approach in the whole healthcare system. Recently, mRNA therapy has gained a great attention as mRNA-based vaccines, e.g. mRNA-1273 (Moderna) [3] and BNT162b2 (Pfizer-BioNTech) [4], became major tools and preferred method of fighting COVID-19 pandemic overworld. However, in addition to long coding sequences, such as pDNA [5], gene-containing viral vectors [6] and mRNA, shorter nucleic acids, i.e. oligonucleotides (ONs), have emerged as highly potent tools in gene therapy. Some of them, e.g. microRNAs (miRNAs) [7], small interfering RNAs (siRNAs) [8] and antisense ONs (ASOs), act by suppressing expression of a harmful gene product, while others, like splice-switching oligonucleotides (SSOs) [9], interact with splice sites on pre-mRNA in order to facilitate synthesis of a normal protein form, release splicing factors or prevent synthesis of aberrant product. SSOs are highly promising novel therapeutic agents as they make possible treatment of genetic disorders that are caused by splicing-impairing mutations in genes, e.g Duchenne muscular dystrophy [10] and spinal muscular atrophy [11]. Some SSOs, such as eteplirsen [10] and nusinersen [11], reached clinics in 2016 and mipomersen [12] in 2013. SSOs that redirect splicing by blocking aberrant splicing-causing sites on pre-mRNA are known as splice-correcting ONs (SCOs) [9] and some of them already are important tools in treatment of certain types of such diseases like X-linked agammaglobulinemia [9] and beta thalassemia [13]. In the current study, we focus on SCO-induced splicing correction, using as a model two cell lines mimicking human beta thalassemia [13, 14] and, accordingly, two different SCOs, targeting different positions on pre-mRNA.

Regardless of the type, the key issue with successful use of ONs in live systems is their insufficient stability and efficiency. Being large and negatively charged molecules, nucleic acids cannot effectively penetrate cell membrane themselves, thus not providing sufficient effect while being introduced into the extracellular environment [15]. Furthermore, unmodified nucleic acids are rapidly degraded in extra- and intracellular compartments, so that transfection efficiency is very low [15, 16]. To address this issue, different chemical modifications are introduced into both sugar and phosphate parts of the sugar-phosphate backbone of oligonucleotides. For example, SCOs used in the present study contain 2’-O-methylation of riboses and substitution of oxygen with sulphur that results in the phosphorothioate backbone, PS, and provides the ON with the greater stability against nucleases as well as higher affinity to the target sequence [17]. However, inefficient cellular delivery of ONs still remains a major issue. Chemical transfection systems used for nucleic acid delivery include a large variety of vehicles [18-20] that aim in increasing the stability of nucleic acid, facilitating its translocation across the plasma membrane, as well as avoiding endosomal entrapment and subsequent degradation of the ON. Among them, cell-penetrating peptides (CPPs) have gained substantial attention as transfection reagents due to their simplicity, high efficiency and low cytotoxicity [21-25].

CPPs are <30 amino acid residues long peptides that can deliver into cells various cargos, such as ONs, pDNA, siRNA, proteins etc. [26] CPPs can be cationic, amphipathic or hydrophobic [20] and some of them form nanoparticles with their cargo molecules that are directly transferred through plasma membrane or enter cells via endocytic pathways [27]. The present study focuses mostly on PepFect14 (PF14), a stearylated amphipathic peptide from the PepFect family. PF14 was first synthesized in 2011 by Ezzat and colleagues [26] and binds its cargos non-covalently, forming nanoparticles that infiltrate into cells mostly by inducing receptor-mediated endocytosis [26, 28, 29]. Members of PepFect family have shown great efficiency in transfecting siRNA [30, 31], miRNA [7, 32-34] and SCO [28, 35], and PF14-SCO nanoparticles have been studied by our group already for some years [28, 29, 36, 37].

Although SCO that is incorporated into PF14-SCO nanoparticles is very effectively transfected into cells and shows lower degradation levels [28] compared with „naked” SCO, novel approaches for increasing efficiency of such nanoparticles are required, especially for delivery *in vivo*. For many transfection systems, endosomal escape is a major bottleneck that determines how large fraction of the molecules can reach their intracellular target. A vector must provide efficient endosomal escape before endosomal pH decreases and hydrolytic enzymes degrade a nucleic acid [38]. Efficiency of endosomal escape can be improved in several ways. For example, membrane fusion can be facilitated by coupling a peptide with fatty acid, like in the case of CPPs from PepFect and NickFect families, and endosome destabilization and escape can be enhanced by incorporating multiple histidine residues into CPP sequence (that results in „proton sponge” effect), as in the case of NF70 and NF72 [39], or by adding endosomolytic residues, like in the case of PepFect6 (PF6) [25]. Still, novel safe strategies are needed to enhance the efficiency of endosomal escape of nanoparticles that carry therapeutic agents.

Based on previous research [37, 40-47], we considered calcium ions to be highly promising candidates for influencing transfection efficiency and biological effect of PF14-SCO nanoparticles. Calcium has been known for long as a compound that is able to condense nucleic acids, thus facilitating transfection of viral DNA [40] and, recently, siRNA [42, 45]. Calcium, as well as magnesium, is known to play multiple roles in cell metabolism, including various aspects of internalization of different substances into cells. For example, it was shown that calcium ions play an important role in endocytosis [47] and receptor transfer to the plasma membrane [36]. In addition, CPP-mediated transfection is accompanied by temporary increase of intracellular calcium concentration [37, 44] and chelation of extracellular calcium ions was shown to reduce both internalization and biological activity of PF14-SCO nanoparticles [37, 44]. It has also been shown that calcium can facilitate transfection of CPP-DNA nanoparticles [41, 43].

Based on it, we decided to test calcium and other biologically relevant divalent metal ions as the constituents of nanoparticles in the cellular models based on splicing correction. Here, we investigate an effect of divalent metal ions, mainly calcium and magnesium, on PF14 nanoparticles with SCO in terms of their physical characteristics, binding to cell surface, transfection efficiency, endosomal escape and splice-correcting activity of SCO. In addition to PF14-SCO nanoparticles, we investigate an effect that calcium and magnesium have on nanoparticles containing several other CPPs and another nucleic acid type, siRNA. We show that, although the effect of calcium and magnesium is not identical for all nanoparticles tested, both substantially increase the biological response to oligonucleotides delivered into cells.

## 2. Materials and methods

### 2.1. Materials

SCO-705 (5’-CCUCUUACCUCAGUUACA-3’), SCO-654 (5’-GCUAUUACCUAAACCCAG-3’) and Cy5-labeled SCO-705 (SCO-705-Cy5) were obtained from Microsynth (Switzerland) and Metabion (Germany). siRNA against luciferase (5’-GGACGAGGACGAGCACUUCTT-3’) and negative siRNA (5’-AGGUAGUGUAAUCGCCUUGTT-3’) were purchased from Microsynth (Switzerland). Following CPPs were used: PF14, NF55, PF6, NF70 (H52), NF71 (H31), NF72 (H82) and C22-PF14 (see STable 1). PF14 was obtained from PepScan (Netherlands), C22-PF14 from Pepmic (China) and all other CPPs were synthesized by Prof. Langel’s group in Stockholm University or Tartu University. 1 mM peptide stock solutions were prepared and stored at -20 °C. Class A scavenger receptor (SR-A) inhibitors polyinosinic acid (Poly I) and fucoidan and their analogues polycytidylic acid (poly C) and galactose were obtained from Sigma-Aldrich (USA, MO). Lipofectamine RNAiMAX Reagent were purchased from Invitrogen (USA, CA). Luciferase substrate XenoLight D-Luciferin – K+ salt (LH2) was obtained from PerkinElmer (USA, MA). Draq5 staining solution was obtained from BioStatus Limited (UK). Other reagents, such as CaCl_2_, MgCl_2_, MnSO4, SDS, chloroquine etc., were obtained from Sigma-Aldrich (Merck KGaA, Germany) and Applichem (USA, IL) if not stated otherwise. 5x Cell Culture Lysis Reagent was obtained from Promega (USA, WI). Ready-to-use Cell Proliferation Colorimetric Reagent, WST-1 was obtained from BioVision (USA, CA). Luciferase activity measurement protocol was based on Helmfors *et al*. 2015 [48] and Rocha *et al*. 2016 [49] and analysis of luciferase activity dependency on different nanoparticle preparation schemes was based on Saher *et al*. 2018 [50].

### 2.2. Cell cultivation

Two human beta-thalassemia reporter cell lines – HeLa pLuc 705 [13] and HeLa EGFP 654 [14, 51] – were used. For siRNA assay, we used luciferase-expressing cell line – U-87 MG-Luc2 [52]. Cells were cultured at 37 °C in Dulbecco’s Modified Eagle Medium (DMEM) (BioWhittaker, USA, or Sigma-Aldrich, UK) that contained 4.5 g/L of glucose and was supplemented with 10% (v:v) of fetal bovine serum (FBS) and 1% (v:v) of penicillin-streptomycin solution (100 U/mL penicillin and 100 μg/mL streptomycin) (further “growth medium”). Cells were grown on 10 cm cell culture plates (Corning, USA, NY) in the humid atmosphere that contained 5% of CO2 and were split every second day. For detaching cells, trypsin-EDTA solution was used (Corning, USA, NY). Cells were washed with Dulbecco’s Phosphate-Buffered Saline (DPBS) without calcium and magnesium (Corning, USA, NY). For experiments, cells were plated on 12-well (Greiner Bio-One, Germany), 24-well (VWR, USA, PA) or 96-well (Sarstedt, Germany) plates. All the operations with cells were performed in sterile cell culture hood.

### 2.3. Nanoparticle formation

For formation of nanoparticles, CPP and SCO were mixed at molar ratio (MR) 5 in Milli-Q (MQ) water in 1/10th of final volume (i.e. 10x nanoparticle solutions). For that, each of reagents was diluted in 1/2 volume of MQ and resulting solutions were blent together (with mixing on vortex for ca 5 seconds). After 15 min incubation at room temperature (RT), CaCl_2_ or MgCl_2_ solutions were added if needed. After 15 min, solutions were diluted with pre-warmed growth medium 10-fold to reach the final volume, and applied to the cells. SCO was applied at a final concentration of 100 nM.

In addition to PF14, other CPPs listed in STable 1 were used for transfection of SCO. Usually, MR5 (CPP:SCO) was used, but MRs of 1, 3, 5, 7, 10, 15 and 20 were also tested. Among bivalent metal ions studied were Ca^2+^ and Mg^2+^ (applied at concentrations from 10 μM to 20 mM) as well as Zn^2+^, Mn^2+^, Fe^2+^ and Cu^2+^ (applied at concentrations from 100 μM to 5 mM). In all experiments, a growth medium containing 10% (v:v) of MQ water was used as a negative control solution.

As an alternative to SCO, siRNA against luciferase was used. It was applied at the concentrations from 5 to 100 nM and for transfection with PF14, molar ratio 17:1 (PF14:siRNA) was used. As a negative control, negative siRNA was used. As a positive control, we made nanoparticles using Lipofectamine RNAiMAX Reagent, according to the manufacturer’s protocol. PF14-siRNA nanoparticles were prepared in the same way as PF14-SCO nanoparticles, diluted 10 times with growth medium and incubated with cells for 48 h before luminescence measurement.

### 2.4. Confocal microscopy

HeLa pLuc 705 cells were plated in a 24-well plate on cover glasses with a diameter of 12 mm (Menzel-Gläser, Germany) 24 h before transfection. Next day, the growth medium was replaced with a nanoparticle-containing medium. For nanoparticle formation, SCO-Cy5 was used, and calcium and magnesium ions were added at concentrations of 0.3 or 3 mM. Cells were incubated with the solutions for 1, 4 or 24 h at 37 °C. Next, cells were washed with DPBS twice and fixed by incubation with 4% paraformaldehyde (PFA) for 30 min at RT. Cell nuclei were stained with 4’,6-diamidino-2-phenylindole (DAPI) 0.5 μg/ml solution in DPBS. Cells were washed with DPBS and mounted to glass slides in 30% glycerol. Specimens were analyzed using Olympus FluoView FV1000 (Olympus, Japan) confocal microscope. For each solution, two specimens were prepared and at least three images per specimen were obtained (ca 10 planes per image with step size of 1.2 μm). Cy5 (λex/em = 633/666 nm) and DAPI (λex/em = 405/470 nm) signals were analyzed and 100x objective with oil immersion was used.

HeLa EGFP 654 cells were plated in chambers (Lan-Tek Chambered #1.0 Borosilicate Coverglass System; Thermo Fisher Scientific, USA, MA). Cells were transfected as described above, using SCO-654. To stain nuclei, cells were incubated in cell culture medium containing 10 μM Draq5 DR50200 reagent (BioStatus, UK) for 3 min at 37 °C. After incubation, cells were washed with DPBS and live cells were analyzed with Olympus FluoView FV1000 microscope. EGFP (λex/em = 488/507 nm) and Draq5 (λex/em = 647/681 nm) signals were analyzed and a 60x objective with water immersion was used. For image editing, FV10-ASW 4.2 Viewer was used. Cy5 and EGFP signal intensity and distribution in cells were analyzed visually and quantitatively, using AutoQuant X3 (Media Cybernetics, USA, MD) and Imaris x64 7.6.5 (Oxford Instruments, UK) softwares.

### 2.5. Electron microscopy of nanoparticles

For analysis of nanoparticle localization and intracellular trafficking in more detail, cells were incubated with nanoparticles assembled with nanogold-labeled SCO (SCO-NG). SCO was labeled with Nanogold and the specimens prepared as described earlier [53]. Briefly, the thiol group at 5′ end of ON was tagged with nanogold cluster (Monomaleimido Nanogold, Nanoprobes, NY, d = 1.4 nm) by forming a covalent bond between the thiol group of oligonucleotide and the maleimide group of label. The non-covalent complexes of SCO-NG with PF14 were formed using the same protocol as used for splice correction experiments described above. HeLa pLuc 705 cells were incubated with complexes of 100 nM SCO-NG with 500 nM PF14 supplemented with 3 mM Ca ions for 4 or 24 h. Cells were then fixed and the nanogold label was magnified by silver enhancement followed by stabilization with 0.05% gold chloride [54]. After post-fixation with osmium tetraoxide, the cells were dehydrated and embedded in epoxy resin. Ultrathin sections were cut in parallel with the coverslip, and post-stained with uranyl acetate and lead citrate. The specimens were examined at 120 kV accelerating voltage with FEI Tecnai G2 Spirit transmission electron microscope (FEI, The Netherlands) and images captured with Orius SC1000 camera.

### 2.6. Flow cytometry

Cells were plated on 12-well plate 24 h prior to transfection. Next day, the growth medium was replaced with nanoparticle-containing medium. Nanoparticles of PF14 and SCO were prepared as described above. For HeLa pLuc 705 cells, Cy5-labeled SCO was used, and for HeLa EGFP 654, SCO-654 was used. Cells were incubated with the nanoparticles for 24 h at 37 °C. Cells were washed with DPBS and collected into 1.5 mL tubes, pulled down and washed with DPBS by centrifugation at 250 rcf at RT. After that, flow cytometry of cell suspensions was performed with BD FACSAria Cell Sorter (BD Biosciences, USA, NJ) and FACSDiva software (BD Biosciences, Germany), using Cy5 or EGFP channel, according to cell line. For each sample, 30 000 events were analyzed.

### 2.7. Luminescence measurement

HeLa pLuc 705 cells were plated on a 96-well plate 24 h prior to transfection. Nanoparticles were prepared and cells were transfected as described above. After incubation, cells were washed twice with DPBS and lysed by adding 20 μL of Cell Culture Lysis Reagent and incubating samples at -20 °C until freezing of the solutions. Then, solutions were thawed at RT, 100 μL of luciferase substrate solution per well was added, solutions were transferred to white 96-well plate (Greiner Bio-One, Germany) and luminescence intensity was measured, using Infinite M200 PRO microplate reader (Tecan, Switzerland) and Magellan 7 software.

### 2.8. Assessment of the role of endosomal escape and SR-As in efficiency of PF14-SCO nanoparticles

To evaluate the efficiency of endosomal escape of differently prepared nanoparticles, we used endosome-destabilizing compound chloroquine [55-57]. Cells were plated on a 96-well plate 24 h prior to transfection. Next day, the growth medium was replaced with a medium that contained nanoparticles and 100 μM chloroquine or nanoparticles only and cells were incubated with these solutions for 4 h. After that, solutions were replaced with fresh growth medium and luciferase activity was measured after additional 20 h of incubation.

To evaluate the role of class A scavenger receptors (SR-As) in transfection and biological efficiency of differently prepared nanoparticles [28, 29], SR-A inhibitors Poly I [58-60] and fucoidan [61-63] were used. Their chemical analogues without inhibiting properties Poly C and galactose, respectively, were used as a negative control. Poly I and Poly C were applied at concentrations of 5, 10 or 25 μg/ml and fucoidan and galactose at concentrations of 2.5 or 5 μg/ml. Cells were plated on a 96-well plate and incubated at 37 °C. Next day, the growth medium was replaced with a medium that contained SR-A inhibitors or their analogues (in 90% of final volume) and cells were incubated with these solutions for 1 h. Then, 10x nanoparticle solutions (in 10% of final volume) were added and luciferase activity was measured after 24 h of incubation.

### 2.9. PCR analysis

HeLa pLuc 705 cells were seeded on a 12-well plate (80 000 cells per well). Next day, SCO-PF14 nanoparticles (100 nM SCO, 500 nM PF14) were prepared with addition of calcium and magnesium or without. Nanoparticles were diluted 10 times with cell culture medium and cells were incubated with the solutions for 24 hours. Next day, cells were harvested and total RNA was extracted using the RNeasy Mini Kit (Qiagen). RNA was subjected to DNase treatment (TURBO DNase, Thermo Scientific) and cDNA was synthesized from 1.34 µg of RNA using RevertAid First Strand cDNA Synthesis Kit (Thermo Scientific). PCR of luciferase gene and GAPDH gene (used as a housekeeper) was performed using Platinum II Hot-Start Green PCR Master Mix (Invitrogen) as described earlier [64]. Agarose gel electrophoresis was performed to analyze the splicing correction, and resulting bands were quantified using ImageJ software.

### 2.10. Nanoparticle cytotoxicity assessment

In aim to assess cytotoxicity of nanoparticles, cell viability was measured spectrophotometrically, using WST-1 assay. Cells were plated on a 96-well plate and transfected the next day as described above. After 24 h of incubation with the solutions studied, 7 μL of WST-1 solution was added to each well and cells were incubated at 37 °C for 3 h. Absorbance of resulting solutions was measured at 440 nm with reference wavelength 650 nm, using Infinite M200 PRO (Tecan) microplate reader. Absorbance of untreated cells was taken for 100% viability.

### 2.11. Measurement of nanoparticle size and zeta potential

Nanoparticle 10x-water solutions were prepared as described above, using 1 μM SCO, 5 μM PF14 and varying concentrations of calcium and magnesium. Solutions were incubated for total of 30 min at RT. For each measurement, nanoparticle 10x solution was diluted 10-fold by adding a solution containing 1 mM MES buffer (pH 7.2) and 1 mM NaCl to reach the final volume of 1 mL. Nanoparticle diameter and zeta potential were measured using ZetasizerNano ZS apparatus (Malvern Instruments, UK).

### 2.12. Statistical analysis

All statistical analyses were performed using Prism 5.0 software (GraphPad Software). Each dataset is presented as the mean ± SEM. Statistical significance of differences between the values of each study group or condition was analyzed using one-way ANOVA with post-hoc Tukey’s or Dunnett’s test at a significance level of 0.05.

## 3. Results and discussion

### 3.1. Addition of divalent metal ions enhances splice-correcting activity of SCO in HeLa pLuc 705 reporter cells

Calcium was known for long as a compound that is able to condense nucleic acids, facilitating transfection of viral DNA [40]. It was previously shown that Ca^2+^ depletion in extracellular environment reduces internalization and biological efficiency of SCO-PF14 nanoparticles [28, 36, 44] and that increased concentration of extracellular calcium has a positive effect on transfection of ONs [46]. Calcium ions have been used to condense siRNA to nanoparticles [45] as well as to facilitate CPP-NA nanoparticle transfection [41-43]. Importantly, calcium-aided transfection is reported to be highly efficient and non-toxic. Based on it, we hypothesized that inclusion of Ca^2+^ ions as well as other divalent metal ions might enhance efficiency of SCO-PF14 nanoparticles and decided to study efficiency and mechanism of such transfection in more detail.

First, we decided to test an effect of addition of divalent metal ions to CPP-SCO nanoparticles in luciferase-based reporter system. For that, HeLa pLuc 705 cells were incubated with ion-complemented or non-complemented PF14-SCO-705 nanoparticles or with SCO-705 alone (Fig. 1). For formation of nanoparticles, CPP and SCO were mixed at a molar ratio (MR) 5 in Milli-Q (MQ) water in 1/10th of final volume, and after 15 min incubation, CaCl_2_ or MgCl_2_ solutions were added to preformed complexes. After 15 min, solutions were diluted with pre-warmed growth medium 10-fold to reach the final volume, applied to the cells and incubated for 24 h. SCO was applied at a final concentration of 100 nM. Calcium and magnesium ions were applied at different concentrations in order to determine minimal effective and optimal concentrations of the ions. Addition of other divalent metal ions that are biologically relevant – Zn^2+^, Mn^2+^, Fe^2+^ and Cu^2+^ – to nanoparticles did not result in any positive change in splicing correction, but was apparently toxic for cells at sub millimolar concentrations (SFig. 2). To assess if the positive effect of the ions is also present at the mRNA level, we incubated HeLa pLuc 705 cells with differently prepared nanoparticles for 24 h and performed PCR analysis of luciferase gene expression (Fig. 1C) [64].

**Fig. 1.**
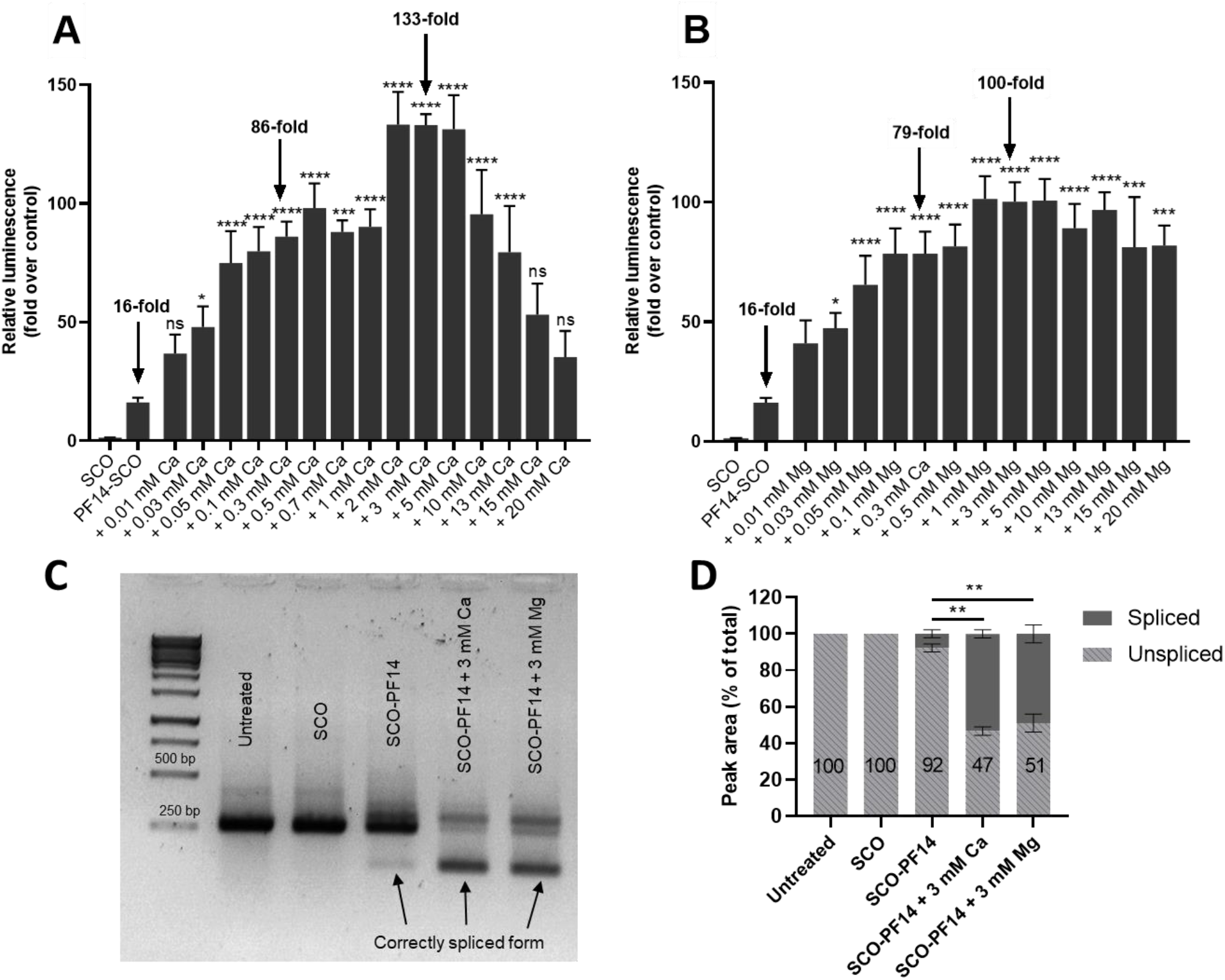
Calcium and magnesium ions increase the biological effect of PF14-SCO nanoparticles in a concentration dependent manner. HeLa pLuc 705 cells were incubated with nanoparticles of 100 nM SCO-705, 500 nM PF14 and with increasing concentrations of Ca^2+^ (A) or Mg^2+^ (B) for 24 h and then luciferase activity was measured. Increase in the proportion of the correctly spliced luciferase mRNA in HeLa pLuc 705 cells as analyzed by PCR (C) and quantification of gel electrophoresis band intensity performed using ImageJ software (D). In the panel C, one representative image is presented. Each dataset in A, B and D represents mean ± SEM of at least 3 independent experiments. Data was analyzed by one-way ANOVA with Dunnett’s test. In the panel D, proportions of aberrantly spliced form of luciferase mRNA are compared. Asterisks indicate statistically significant difference compared to PF14-SCO, *p-value < 0.05, ***p-value < 0.005, ***p-value < 0.0005, ****p-value < 0.0001.

Both calcium and magnesium ions caused significant increase in SCO splice-correcting activity while added to nanoparticles at concentrations of 0.03 mM and higher. However, the effect of calcium and magnesium was different. Positive effect of calcium increased with its concentration, reaching maximum at 2-5 mM, and then decreased dramatically, completely disappearing at 15 mM (Fig. 1A). At the same time, the effect of magnesium neither depended so strongly on its concentration, nor disappeared at high concentrations (Fig. 1B). At concentrations under 10 mM, calcium was more effective than magnesium, resulting in nearly 8-fold increase of biological efficiency of PF14-SCO nanoparticles while applied at concentrations of 2-5 mM. However, at higher concentrations, effect of magnesium was not reduced, making it more efficient, compared to calcium. For the following experiments, we decided to use 0.3 mM calcium, as it produced intermediate, but statistically significant increase (5.3-fold) of luciferase expression, and 3 mM calcium, as it was the optimal concentration for reaching maximal activity and significantly differed from the effect of 0.3 mM calcium (p < 0.0001). Also, we decided to use magnesium at the same concentrations, to make the non-specific effect of the ions comparable. Addition of calcium and magnesium significantly increased the proportion of shorter, correctly spliced form of the mRNA, as verified by quantification of bands (Fig. 1D). The efficiency of “naked” SCO did not change upon addition of divalent metal ions (data not shown).

### 3.2. Adding calcium and magnesium ions to SCO-PF14 nanoparticles yields nanoparticles with similar physical properties and does not impair viability of the cells

In order to assess, whether the addition of calcium and magnesium ions to SCO-PF14 nanoparticles changes their physical properties, we measured size and zeta potential of the nanoparticles. Hydrodynamic diameter and zeta potential were measured with ZetaSizer, using buffer with minimal ionic strength to analyze nanoparticles at concentration used in experiments with cells (Table 1). Our results indicate that PF14 forms particles with diameter about 230 nm with SCO. Adding calcium and magnesium to the nanoparticles increases their diameter to some extent, though not significantly, and no evident concentration dependence is observed. All nanoparticles have hydrodynamic diameter under 500 nm, thus staying in physiologically safe size range. Nanoparticles of SCO and PF14 have a positive zeta potential of about +9.3 mV. Upon addition of calcium, zeta potential did not change markedly, indicating that incorporation of calcium and magnesium ions into the nanoparticles does not substantially influence the charge of PF14-SCO nanoparticles.

**Table 1.** The effect of adding calcium and magnesium ions on biophysical characteristics of PF14-SCO nanoparticles. Nanoparticle 10x water solutions were prepared by mixing 1 µM SCO and 5 µM PF14 in MQ water and adding 3 or 30 mM calcium and magnesium 15 minutes later. The solutions were incubated for total of 30 min at room temperature. For each measurement, nanoparticle 10x solution was diluted 10-fold by adding a solution containing 1 mM MES buffer and 1 mM NaCl to reach the final volume of 1 mL. Nanoparticle diameter and zeta potential were measured using ZetasizerNano. Each dataset represents mean of at least three independent experiments (performed in triplicates) ± SEM. Data was analyzed by one-way ANOVA with Tukey’s test; no significant differences were detected.

| Nanoparticle | Size (nm) | Zeta Potential (mV) |
| --- | --- | --- |
| PF14-SCO | 231±22 | 9.3±1.1 |
| PF14-SCO + 0.3 mM CaCl<sub>2</sub> | 327±60 | 10.1±1.8 |
| PF14-SCO + 3 mM CaCl<sub>2</sub> | 305±51 | 13.1±1.0 |
| PF14-SCO + 0.3 mM MgCl<sub>2</sub> | 352±65 | 10.6±1.7 |
| PF14-SCO + 3 mM MgCl<sub>2</sub> | 393±60 | 15.0±2.4 |

Possible effect of adding calcium ions into SCO-PF14 nanoparticles on viability of the cells was analyzed using WST-1 assay (Fig. 2). HeLa pLuc 705 cells were incubated with nanoparticle-containing solutions for 24 h. Then, WST-1 reagent was added, cells were incubated with it for 3 h at 37 °C and absorbance was measured according to the manufacturer’s protocol. Neither naked SCO nor SCO-PF14 nanoparticles (at any molar ratio from 1 to 20) affected viability of HeLa pLuc 705 cells. Adding 0.1 mM to 10 mM calcium and magnesium to the nanoparticles did not lead to any decrease in the cell viability. Also, adding PF14 or calcium and magnesium alone did not affect viability of the cells, unlike zinc and manganese, that were highly cytotoxic at submillimolar concentrations (SFig. 2).

**Fig. 2.**
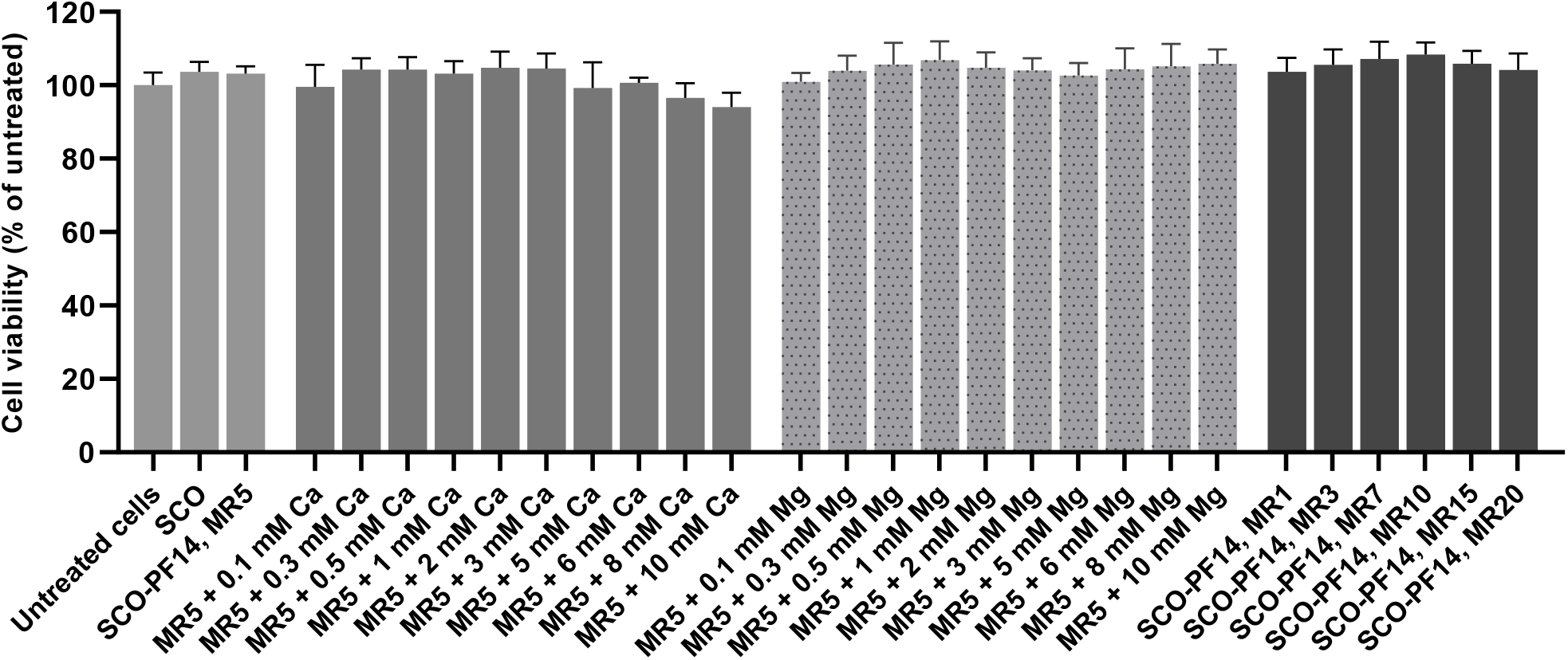
The effect of SCO, PF14, calcium and magnesium ions on viability of cells. HeLa pLuc 705 cells were incubated for 24 h with solutions containing SCO alone (100 nM), nanoparticles of SCO and PF14 taken at different molar ratios (MR) or calcium or magnesium-complemented nanoparticles. Viability of the cells was evaluated using WST-1 assay, and absorption of untreated cells was taken for 100%. Each dataset represents mean + SEM of at least 3 independent experiments. Data was analyzed by one-way ANOVA with Dunnett’s test, no statistically significant differences were detected.

### 3.3. Association of PF14-SCO complexes with cells and intracellular distribution of SCO changes upon addition of divalent metal ions into nanoparticles

In order to shed light on the mechanism of calcium and magnesium-induced increased splicing correction and to study intracellular distribution of nanoparticles, we harnessed SCO labeled with Cy5. HeLa pLuc 705 cells were incubated with ion-complemented or non-complemented PF14-SCO-Cy5 nanoparticles or with SCO-Cy5 alone (Fig. 3), applied at a final concentration of 100 nM. Cells were incubated with the respective solutions for 1, 4 or 24 h to analyze the association and uptake by cells, escape from endosomes and translocation into nucleus, respectively.

**Fig. 3.**
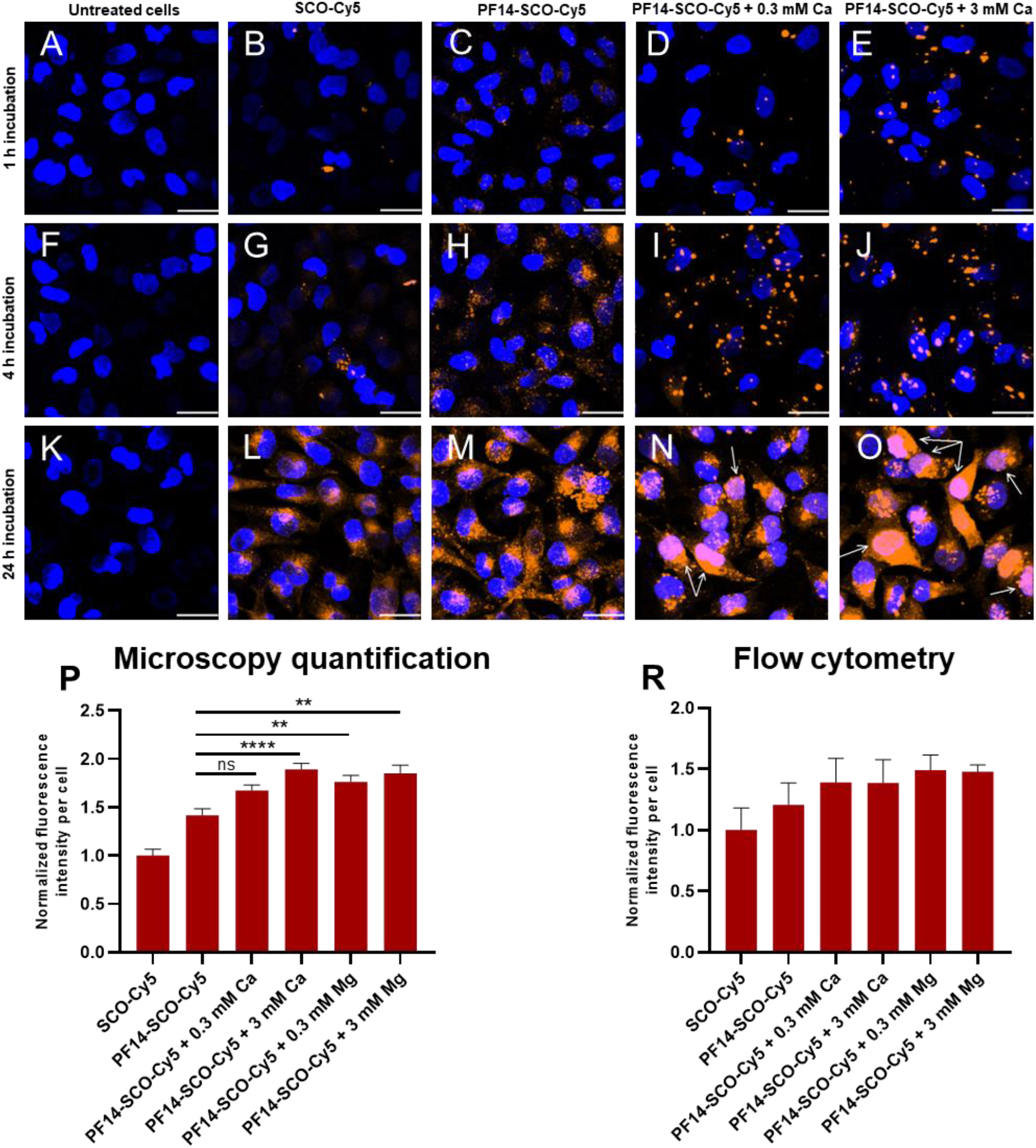
Delivery of splice-correcting oligonucleotide into HeLa cells with PF14 nanoparticles containing calcium ions. HeLa pLuc 705 cells were incubated with nanoparticles of 100 nM SCO-Cy5, 500 nM PF14 and 0.3 or 3 mM Ca^2+^ for 1, 4 or 24 h, fixed and localization of SCO by Cy5 fluorescence (orange signal) was analyzed with Olympus FluoView FV1000 (Olympus, Japan) confocal microscope. Cell nuclei were visualized with DAPI (blue). Inclusion of Ca ions into PF14-SCO nanoparticles led to distribution of SCO-Cy5 to cytosol and nucleus of cells (with arrows on L and O). The merged images of all confocal planes are presented. One representative of three independent experiments is presented. Scale bar 30 μm. Panel P: Cy5 signal of at least 100 cells per sample was quantified using AutoQuant X3 software. Panel R: alternatively, flow cytometry analysis was performed to analyze association and internalization of PF14-SCO-Cy5 nanoparticles after 24 h, using BD FACSAria Cell Sorter (BD Biosciences, USA). In flow cytometry, 30 000 events per sample were detected, each experiment was performed at least 3 times. All values are normalized against average fluorescence intensity in the samples treated with SCO alone. Each dataset represents mean + SEM. Data was analyzed by one-way ANOVA with Tukey’s test, **p-value < 0.005, ****p-value < 0.0001, ns – not significant.

PF14-SCO-Cy5 nanoparticles produced granular signal, that likely represents invaginations and endosomes forming upon internalization of nanoparticles (Fig. 3C, H, M). Importantly, in the case of SCO-PF14 nanoparticles, fluorescence signal was detectable after 1 h incubation (Fig. 3C) and was significantly stronger compared to SCO-Cy5 alone. In the case of “naked” SCO-Cy5, there was a punctuate staining that was rather evenly distributed in cell cytoplasm (Fig. 3B, G, L) and clearly detectable, although very weak, after 4 h of incubation (Fig. 3G). Upon addition of calcium ions, bigger Cy5 signal “granules” formed (Fig. 3D, E, I, J, N, O), that were clearly seen already after 1 h of incubation, and their size and shape varied substantially, suggesting that the PF14-SCO nanoparticles could form agglomerates or be internalized in larger vesicles. After 24 h incubation, in parallel with large vesicles, a diffuse Cy5 signal was also observable in the cells treated with ion-complemented nanoparticles (Fig. 3N, O). Remarkably, after 24 h incubation, Cy5 signal was abundant in cell nuclei in case of specimens treated with ion-complemented nanoparticles (Fig. 3N, O, white arrows), suggesting that addition of calcium and magnesium ions drastically increases the amount of SCO that reaches the cell nucleus, i.e. site of activity. Magnesium ions in NPs produced an effect that was analogous with calcium (not shown).

The effect of calcium and magnesium ions on SCO delivery into cells with PF14 was quantitatively analyzed in the confocal microscopy images (Fig. 3P). For that, Cy5 signal intensity of the samples with 24 h incubation (Fig. 3K-O) was quantified, using AutoQuant X3 software. Signal quantification corroborated significantly, though only up to 34% higher internalization of calcium-complemented nanoparticles into HeLa pLuc 705 cells (Fig. 3P). Flow cytometry gave very similar results (Fig. 3R), showing slight increase (up to 24%, not significant) of Cy5 signal upon complementing SCO-PF14 nanoparticles with calcium and magnesium after 24 h incubation.

Next, we used another reporter cell line, HeLa EGFP 654 [14], in order to examine, whether the positive effect on splice-correction activity of SCO upon addition of calcium and magnesium ions to PF14-SCO nanoparticles is universal, and could be recapitulated with another reporter protein, i.e. EGFP. For that, cells were transfected with ion-complemented or non-complemented PF14-SCO-654 nanoparticles or with SCO-654 alone and EGFP signal was captured in live cells after 24 h incubation (Fig. 4). Confocal microscopy revealed substantial growth of the signal intensity upon addition of calcium and magnesium ions (Fig. 4D, E), indicating higher splice-correcting activity of SCO-654, that resulted in higher expression of EGFP, compared to samples that were treated with nanoparticles containing only PF14 and SCO (Fig. 4C).

**Fig. 4.**
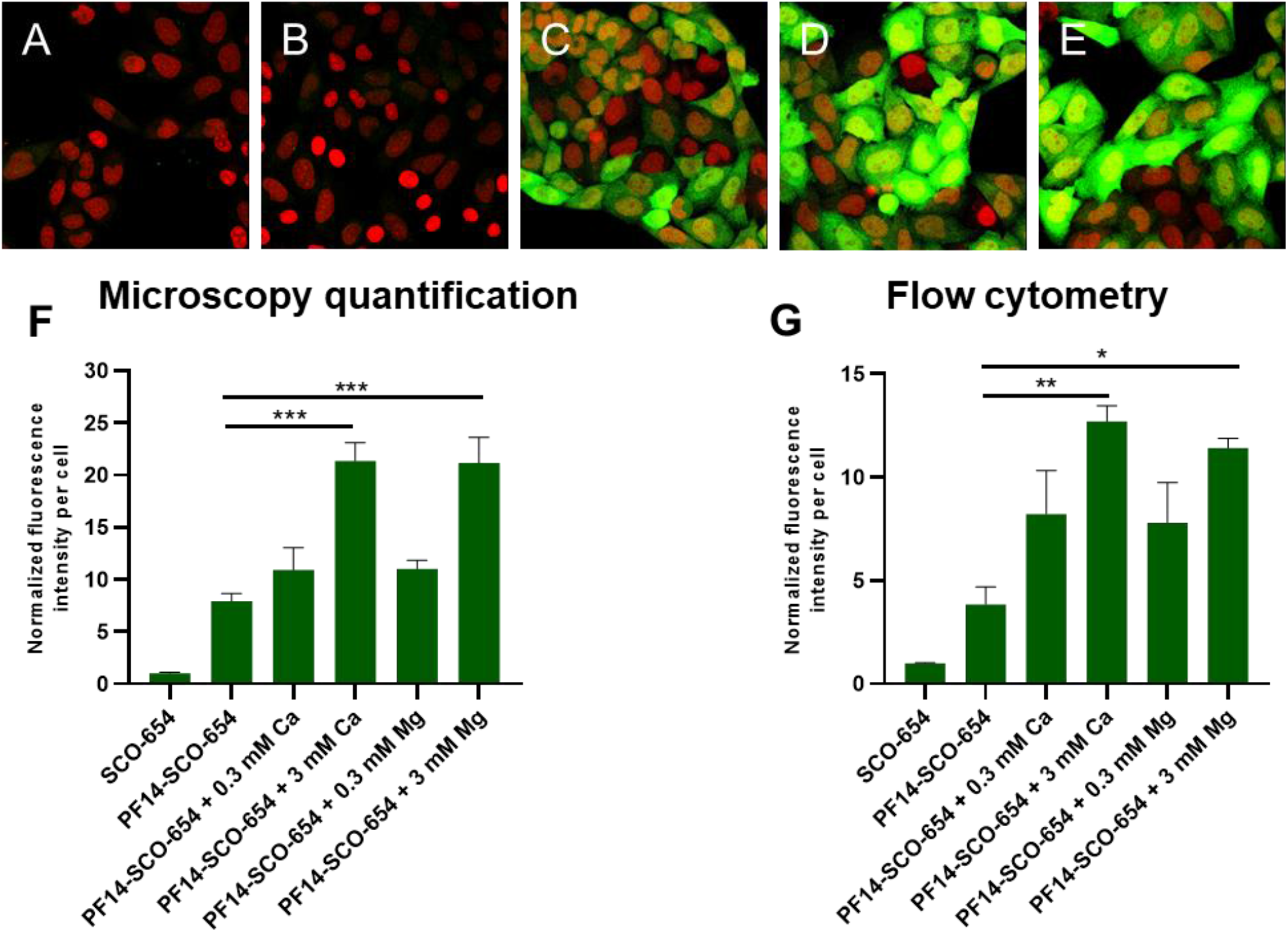
Enhancement of EGFP expression by inclusion of Ca^2+^ or Mg^2+^ ions into PF14-SCO nanoparticles for increasing splicing correction in HeLa EGFP reporter cells. HeLa EGFP 654 cells were incubated with nanoparticles of 100 nM SCO-654, 500 nM PF14 and 3 mM Ca^2+^ or Mg^2+^ for 24 h for triggering splicing switching. Rescued EGFP expression (green) was analyzed with Olympus FluoView FV1000 confocal microscope. Cell nuclei were visualized with Draq5 (red). A, untreated cells; B, cells treated with SCO-654; C, PF14-SCO-654; D, PF14-SCO-654 + 3 mM Ca^2+^; E, PF14-SCO-654 + 3 mM Mg^2+^. The merged images of all confocal planes are presented. One representative of three independent experiments is presented. Scale bar 30 μm. Panel F: EGFP signal of at least 100 cells per sample was quantified using AutoQuant X3 software. Panel G: alternatively, flow cytometry analysis was performed to analyze association and internalization of PF14-SCO-Cy5 nanoparticles after 24 h, using BD FACSAria Cell Sorter (BD Biosciences, USA). In flow cytometry, 30 000 events per sample were detected, each experiment was performed at least 3 times. All values are normalized against average fluorescence intensity in the samples treated with SCO alone. Each dataset represents mean + SEM. Data was analyzed by one-way ANOVA with Tukey’s test, *p-value < 0.05, **p-value < 0.005, ***p-value < 0.0005.

The fluorescence intensity was quantified in the confocal microscopy images (Fig. 4F). It showed markedly higher EGFP expression, i.e. splice-correction activity of SCO-654, upon addition of 3 mM calcium or magnesium to PF14-SCO-654 nanoparticles. Adding 3 mM calcium or magnesium ions led to up to 2.5-fold increase of the efficiency of the nanoparticles. Flow cytometry corroborated these results, also showing significant, up to 3-fold increase of fluorescence intensity upon complementing the nanoparticles with calcium and magnesium ions (Fig. 4G).

Thus, even though the amount of SCO entering cells only slightly increases upon addition of calcium and magnesium ions to nanoparticles, a markedly higher amount of SCO reaches cell nuclei (Fig. 3N, O), that results in significantly higher splice-correction activity of SCO (Fig. 1, 4). Altogether, these results suggest that increased activity of ion-complemented nanoparticles cannot be assigned solely to the increased internalization, but is more likely connected to the changes in their intracellular trafficking and resulting more extensive nuclear localization of SCO that, in turn, facilitates higher biological activity of the oligonucleotide.

Next, transmission electron microscopy (TEM) was harnessed to gain a more detailed insight into association of ion-complemented CPP-SCO nanoparticles with cells and following intracellular trafficking (Fig. 5). For localizing every SCO molecule in TEM images, the nanoparticles were assembled using Nanogold-labeled SCO (SCO-NG) [36], using the concentrations identical with fluorescence microscopy: 100 nM SCO (SCO-NG), 500 nM PF14 and 3 mM Ca^2+^. SCO-NG containing nanoparticles accumulated on the surface of HeLa cells, associating with flat membrane areas and filopodia-like protrusions (Fig. 5A, B, C). The size of SCO-NG containing nanoparticles on cell surface corresponded to size of particles with unmodified SCO measured by DLS (Table 1), suggesting that PF14-SCO-Ca^2+^ particles did not aggregate in serum-containing cell culture medium. The cellular uptake pattern of calcium-containing nanoparticles was reminiscent of PF14-SCO complexes internalization observed earlier: mostly macropinocytosis was induced by such nanoparticles, and clathrin-as well as caveolin-dependent endocytosis was triggered at lower extent [28].

**Fig. 5.**
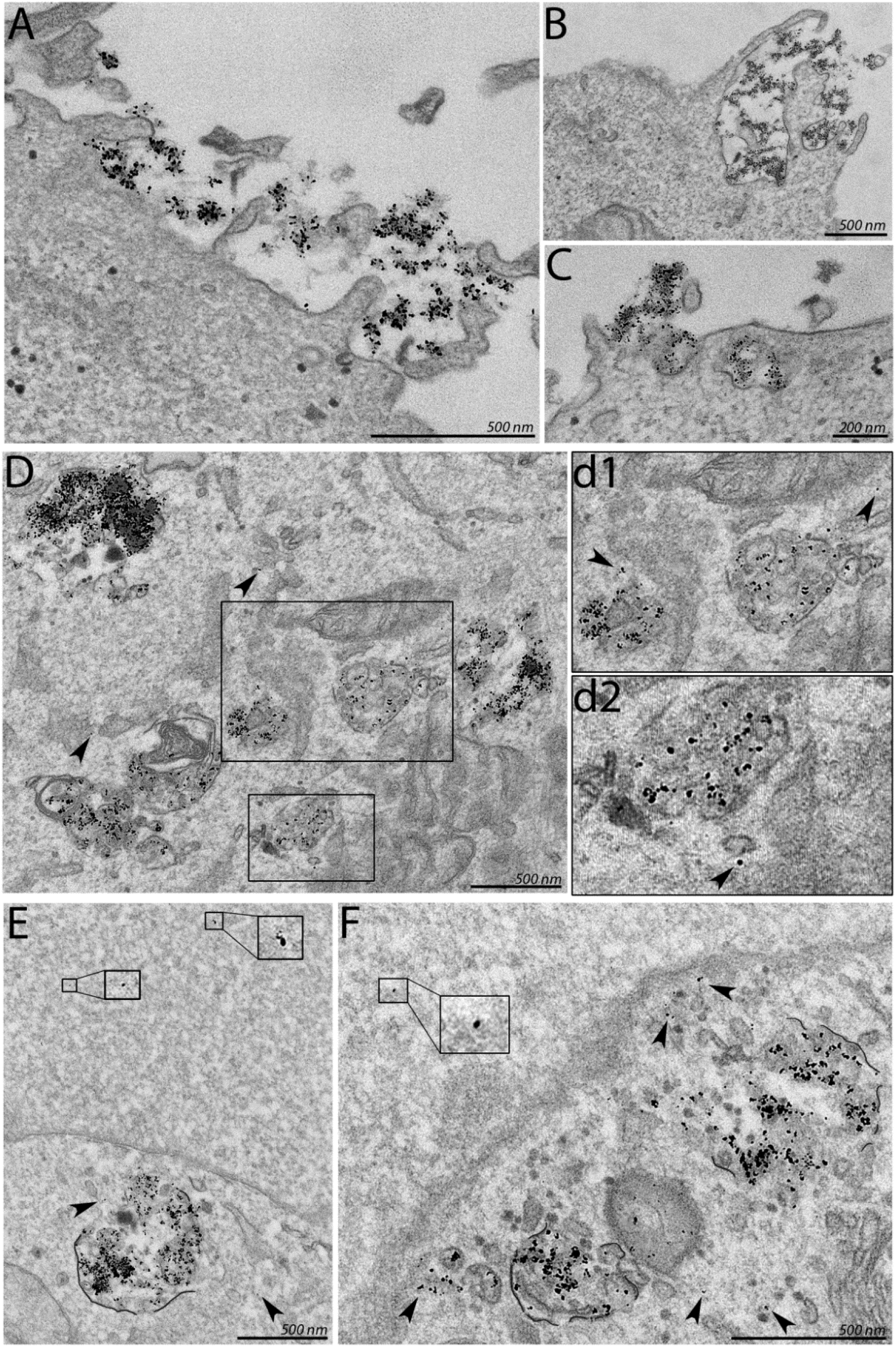
Internalization, endosomal escape and nuclear localization of calcium-complemented CPP-SCO nanoparticles. Cells were incubated with nanoparticles of 100 nM nanogold-labeled SCO (SCO-NG), 500 nM PF14 and 3 mM Ca^2+^ for 4 h (A, B, C) or 24 h (D, E, F) and the specimens were examined at 120 kV accelerating voltage with FEI Tecnai G2 Spirit transmission electron microscope (FEI, The Netherlands). A, association of nanoparticles with plasma membrane. Small black dots represent nanogold label and SCO. B and C, internalization of nanoparticle by cell using endocytosis. D, endosomes containing nanoparticles; d1 and d2, endosomal escape of nanoparticles. E and F, SCO-NG in cell nucleus (magnified areas) and near nuclear membrane (F). Arrowheads indicate SCO-NG in cytosol.

Electron microphotos demonstrate that calcium-containing nanoparticles release SCO from endosomes into cell cytosol more efficiently compared to PF14-SCO nanoparticles not supplemented with divalent metal ions (arrowheads in Fig. 5D, d1, d2), especially in the perinuclear area (Fig. 5E, F). Importantly, within 24 hours, SCO was transported into cell nucleus, i.e. site of activity in easily detectable amounts, notwithstanding the Nanogold label. Both, electron and fluorescence microscopy indicate that Ca^2+^ and Mg^2+^ ions included into PF14-SCO nanoparticles contribute more into intracellular trafficking and redistribution of oligonucleotide than to the uptake of particles by cells.

### 3.4. Calcium ions increase the efficiency of different CPPs

Inspired by the strong positive effect of calcium/magnesium ions on PF14-mediated cellular delivery of SCO, we also tested other cell-penetrating peptides of PepFect and NickFect families: PF6 [25], C22-PF14 [22], NF55 [65], NF70 (H52) [39], NF71 (H31) [39] and NF72 (H82) [39] (Fig. 6). Nanoparticles were formed as described above for PF14 from 100 nM SCO-705 and 500 nM CPP, and luciferase activity was measured after 24-h incubation with HeLa pLuc 705 cells. NF70 and NF72, that were designed for siRNA delivery, were rather inefficient at the transfection of SCO, increasing luminescence less than 6-fold, compared to untreated control. Nanoparticles formulated with PF6 were more efficient in triggering splicing correction by SCO, although their efficacy was not influenced by the presence of calcium ions, producing about 30-fold increase of luminescence of the cells. The efficiency of NF71, NF55 and C22-PF14 at transfecting SCO was strongly influenced by addition of calcium, similarly to PF14. While NF71 showed slightly lower efficiency, NF55 and C22-PF14 were comparable to PF14 at increasing bioavailability of SCO in cells.

**Fig. 6.**
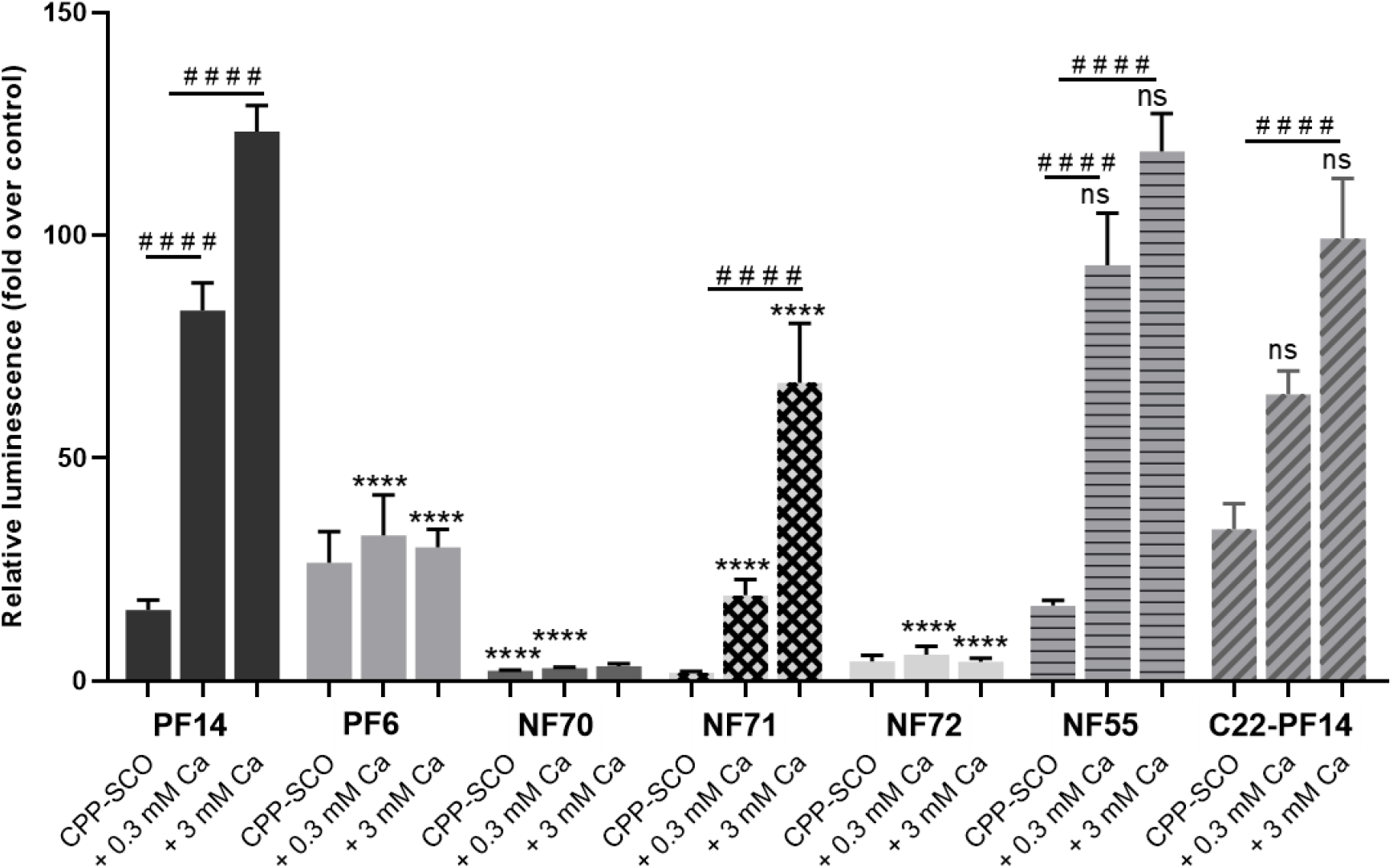
Effect of calcium ions on biological efficiency of nanoparticles formed with different cell-penetrating peptides. HeLa pLuc 705 cells were incubated with nanoparticles of 100 nM SCO-705, 500 nM CPP and 0.3 or 3 mM Ca^2+^ for 24 h, and luciferase activity was measured. Each dataset represents mean + SEM of at least 3 independent experiments. Data was analyzed by one-way ANOVA with Dunnett’s test. Asterisks indicate statistically significant difference compared to corresponding nanoparticle with PF14 (i.e. with one of first three columns), ****p-value < 0.0001, ns – not significant. Hashes indicate statistically significant difference within one group, ####p-value < 0.0001.

### 3.5. Calcium ions reduce the effect of endosome-destabilizing agent chloroquine on splicing correction in concentration-dependent manner

To investigate whether endosomal trapping is a significant hindrance also for activity of ion-complemented PF14-SCO nanoparticles, endosome-destabilizing compound chloroquine was used (Fig. 7). HeLa pLuc 705 cells were incubated with nanoparticles and 100 μM chloroquine-containing growth medium for 4 h. After that, the solutions were exchanged to fresh growth medium and luciferase activity was measured after 20 h. In line with what was expected, chloroquine enhanced efficiency of PF14-SCO nanoparticles (over 9-fold increase of luminescence). However, positive effect of chloroquine decreased at higher calcium ion concentrations, being 3.8-fold in case of 0.3 mM calcium and only 1.5-fold in case of 3 mM calcium. Thus, it can be concluded that calcium-complemented nanoparticles are less dependent on additional endosome destabilizing, i.e. these are more successful at endosomal escape.

**Fig. 7.**
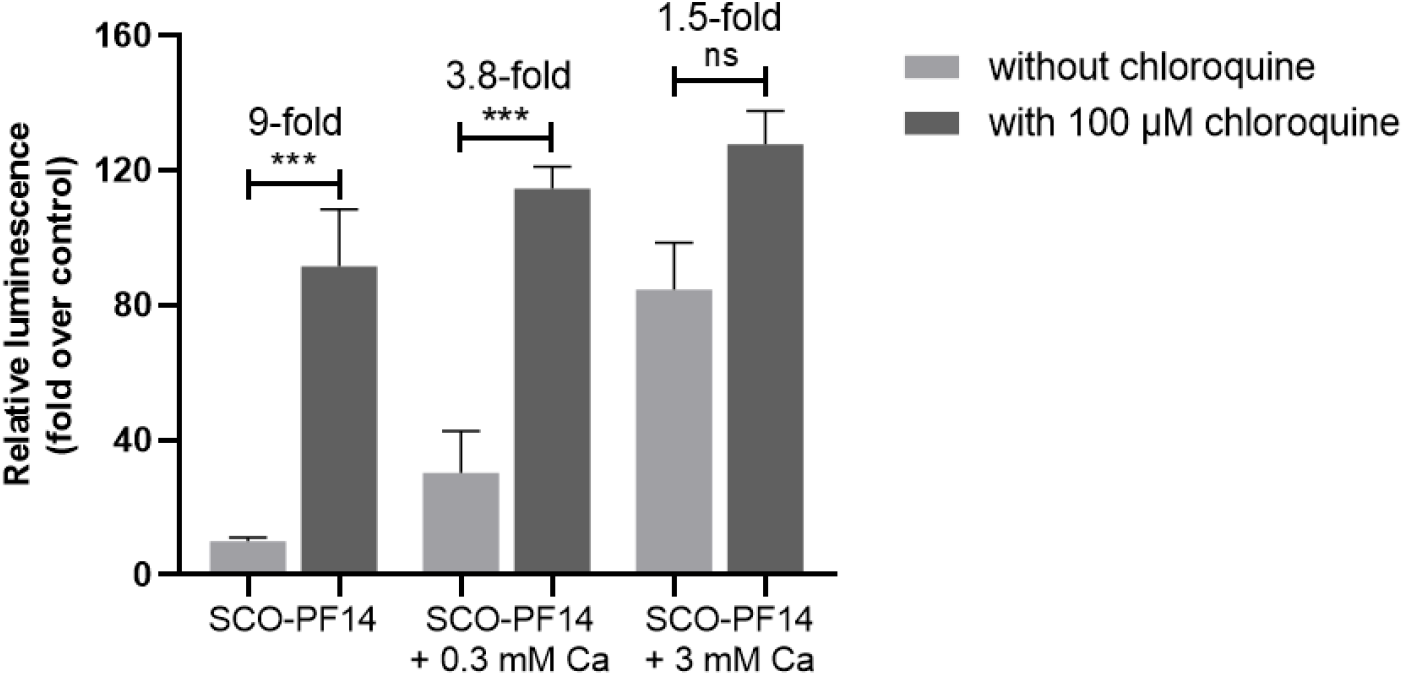
SCO-PF14 nanoparticles rescue luciferase expression in HeLa pLuc 705 cells and reduce effect of endosomolytic agent chloroquine. HeLa pLuc 705 cells were incubated with nanoparticles of 100 nM SCO, 500 nM PF14 and 0.3 or 3 mM Ca^2+^ in growth medium with or without 100 μM chloroquine. After 4 h, the solutions were exchanged to fresh growth medium and luciferase activity was measured after 20 h. Each dataset represents mean + SEM of 3 independent experiments. Data was analyzed by one-way ANOVA with Tukey’s test. Asterisks indicate statistically significant difference between two datasets, ***p-value < 0.0005, ns – not significant.

### 3.6. SR-A inhibitors reduce biological activity of ion-complemented nanoparticles in concentration-dependent manner

We have shown earlier that for PF14-SCO nanoparticles, the main route for entering cells is receptor-mediated endocytosis and one class of receptors required for their entrance are class A scavenger receptor (SR-As), namely SR-A3 and SR-A5 [28, 29]. To find out if the main route of entering cells for ion-complemented PF14-SCO nanoparticles is also SR-A-mediated endocytosis, the receptors on HeLa pLuc 705 cells were blocked with their natural ligands – Poly I and fucoidan. As a negative control, chemical analogues of these ligands were used – Poly C and galactose, respectively. Receptor inhibitors or their analogues were added to HeLa pLuc 705 cells one hour before transfection with nanoparticles. Then, PF14-SCO nanoparticles were added and luminescence was measured after 24 h (Fig. 8).

**Fig. 8.**
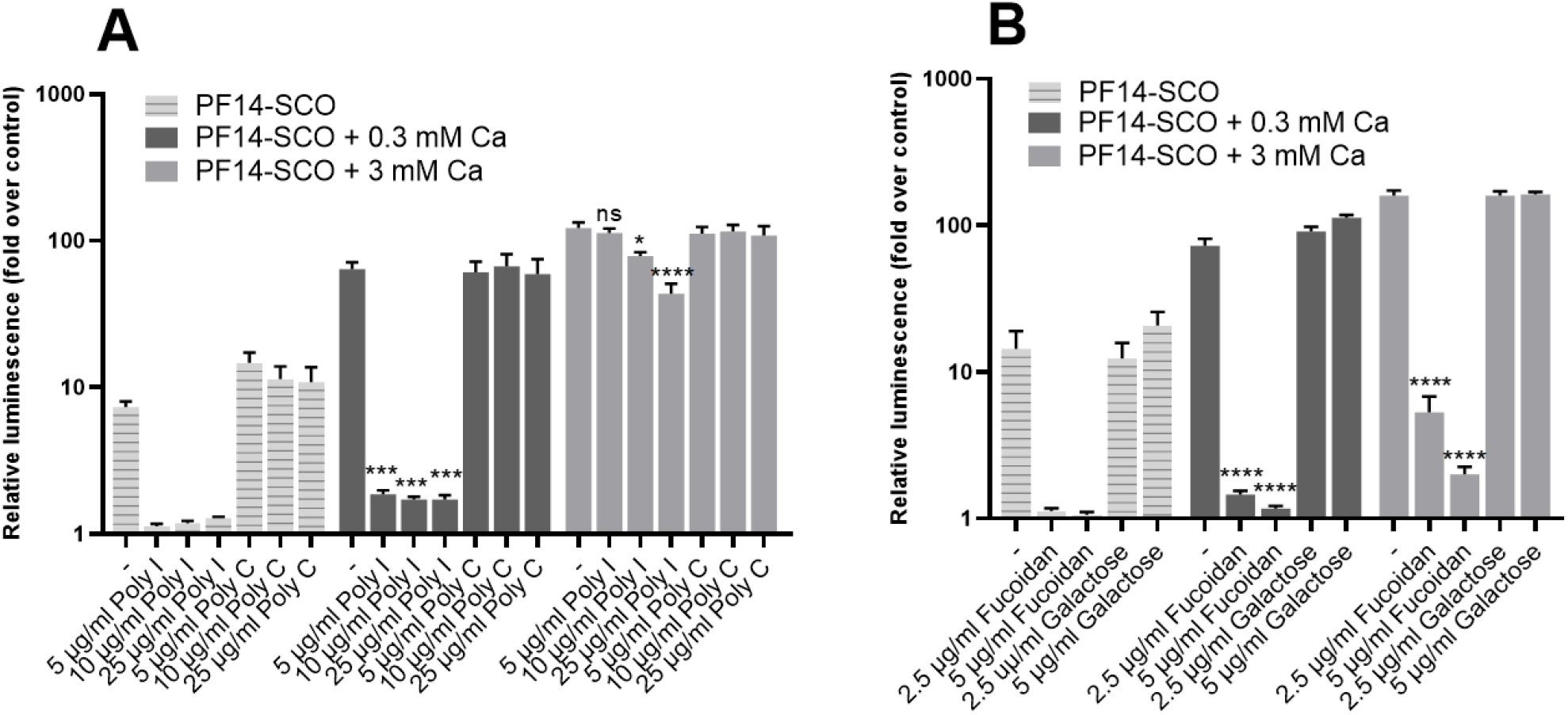
The effect of inhibitors of class A scavenger receptors, SR-As, on biological activity of PF14-SCO nanoparticles. HeLa pLuc 705 cells were incubated with solutions containing SR-A inhibitors Poly I (A) or fucoidan (B) that were added to growth medium at different concentrations. As a negative control, chemicals with analogous structure to the inhibiors, Poly C and galactose, respectively, were used. After 1 h incubation, nanoparticles of 100 nM SCO, 500 nM PF14 and 0.3 or 3 mM Ca^2+^ were added and luminescence was measured after 24 h. Each dataset represents mean + SEM of 3 independent experiments. Data was analyzed by one-way ANOVA with Tukey’s test. Asterisks indicate statistically significant difference with the same nanoparticle without inhibitor or its analogue added, *p-value < 0.05, ***p-value < 0.0005, ****p-value < 0.0001, ns – not significant.

Results of experiments with SR-A inhibitors showed, in concordance with earlier data, that SR-As are essential for endocytosis of non-complemented SCO-PF14 nanoparticles, as their blockage leads to reduction of luminescence to near-basal level. However, blockage of SR-As had less prominent effect when calcium ions were added into nanoparticles, and Poly I at highest, 25 ug/ml, concentration led only to a 37% reduction of luminescence in the presence of 3 mM calcium (Fig. 8A). That may indicate that ion-complemented nanoparticles compete more efficiently with natural ligands of scavenger receptors, or that alternative routes of internalization are used by these nanoparticles.

### 3.7. Addition of divalent metal ions enhances RNA interference effect of siRNA-PF14 nanoparticles in U87-luc2 reporter cells

Encouraged by the strong positive effect of calcium and magnesium ions on the bioavailability of SCO, we analyzed the potentiation of an alternative short nucleic acid. For that, we used luciferase-suppressing siRNA (Fig. 9). We applied it at the concentrations from 5 to 100 nM and for transfection with PF14, we used molar ratio 17:1 (PF14:siRNA), that corresponds to the same N:P (charge) ratio as used in PF14-SCO nanoparticles. As a negative control, we used negative siRNA, and as a positive control, we made nanoparticles using Lipofectamine RNAiMAX Reagent (Invitrogen, CA, USA). PF14-siRNA nanoparticles were prepared in the same way as for SCO, diluted 10 times with growth medium and incubated for 48 h before luminescence measurement.

**Fig. 9.**
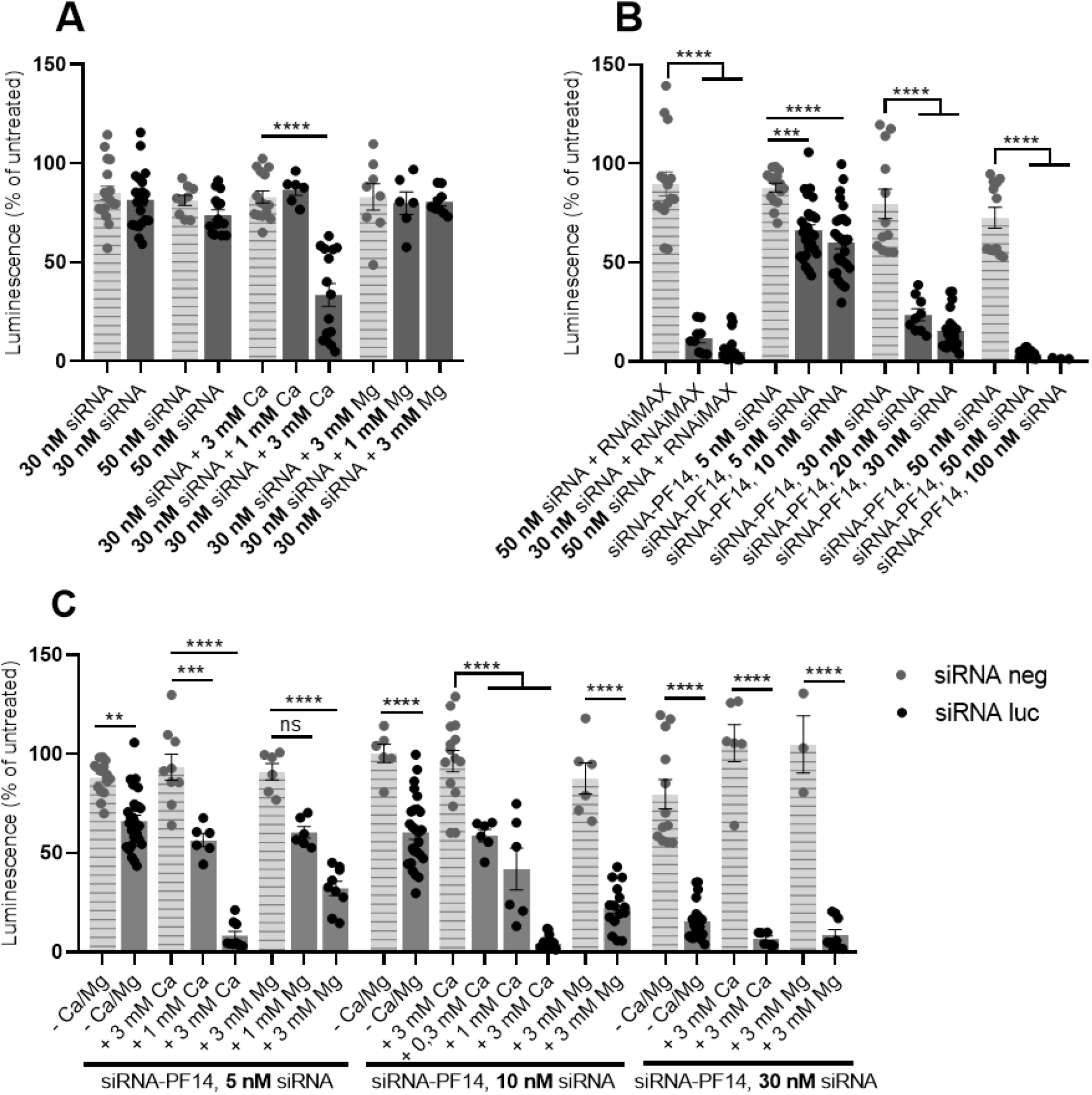
Effect of calcium and magnesium ions on gene siRNA-induced silencing. U87-luc2 cells were incubated with nanoparticles of 5 to 100 nM siRNA, PF14 or Lipofectamine RNAiMAX (positive control) and 0.3 to 3 mM Ca^2+^ or Mg^2+^. For siRNA-CPP nanoparticle preparation, molar ratio 17 was used (PF14:siRNA), that corresponds to charge (N:P) ratio of 2. After 48 hours of incubation, luciferase activity was measured. As a control, negative siRNA was used. Each dataset represents mean ± SEM of at least 3 independent experiments. Luminescence of untreated cells is taken for 100%. Data was analyzed by one-way ANOVA with Tukey’s test, and asterisks indicate statistically significant difference between indicated datasets, **p-value < 0.005, ****p-value < 0.0005, ****p-value < 0.0001, ns – not significant.

PepFect14 turned out to be as good vehicle for siRNA transfection as RNAiMAX: both provided about 95% decrease of luciferase expression in the case of 50 nM siRNA, compared to the analogous nanoparticles that contained negative siRNA (Fig. 9B). Next, we tested an addition of calcium and magnesium to „naked” siRNA (Fig. 9A) and to the siRNA-PF14 nanoparticles (Fig. 9C). In concordance with earlier research, calcium significantly improved efficiency of “naked” siRNA [45], providing (at 3 mM) 60% decrease of the gene expression, compared to the negative siRNA-containing nanoparticles (Fig. 9A). At the same time, magnesium, applied at the same concentration, did not affect the efficiency of siRNA (Fig. 9A). However, both ions significantly improved efficiency of siRNA-PF14 nanoparticles (Fig. 9C), although the effect of calcium was generally stronger than the effect of magnesium applied at the same concentration. Addition of ions produced more efficient nanoparticles in the case of all siRNA concentrations tested. Interestingly, 1 mM calcium did not have any effect on the „naked” siRNA transfection (Fig. 9A), but in the case of siRNA-PF14 nanoparticles, addition of 0.3 mM calcium was sufficient to see a significant effect (Fig. 9C). It indicates that higher amounts of calcium ions are required to improve efficiency of siRNA itself than of siRNA-PF14 complexes. That may be due to stronger negative charge of the former, that is already partially neutralized by the positively charged peptide in the case of the PF14-siRNA nanoparticles. In addition, differing effects of calcium and magnesium give reason to think that there are substantial differences in the mechanisms underlying increased efficiency of the nanoparticles complemented by different ions, that may become apparent only in the case of some types of cargos.

## 4. Discussion

Cell-penetrating peptides (CPPs) are promising tools for nucleic acid (NA) transfection. They are efficient and non-toxic vectors for delivery of various types of oligonucleotides with biological activity, e.g. siRNA, miRNA and SCO. Furthermore, CPPs can also be used to transport into cells nucleic acids with high molecular mass, like pDNA and mRNA, and other bioactive macromolecules, like peptides, proteins, as well as small molecules [66-68]. However, they are still often less efficient than analogous lipid-based delivery systems. One obstacle for efficient use of CPPs is endosomal entrapment of CPP-NA nanoparticles and subsequent degradation of bioactive cargo. Efficiency of endosomal escape can be improved in several ways, for example by using the „proton sponge” effect or by adding endosomolytic residues to the CPP, like in the case of PepFect6 (PF6) [25]. However, CPP-based transfection still needs new ways of improvement, as well as deeper understanding of mechanisms underlying nanoparticle binding to cell surface, translocation through the plasma membrane and/or endocytosis, intracellular trafficking, endosomal escape and binding to intracellular target.

Previously, we showed that CPPs split into two different groups based on intracellular calcium concentration change that follows their entry into cells [37]. Transient increase of intracellular calcium concentration was also shown by other researchers [44] and seems to be an important step for CPP-facilitated transfection. In line with this, it was shown that the effect of oligonucleotides can be enhanced by increasing extracellular calcium concentration [46]. Calcium was known for long as a compound that is able to condense nucleic acids, facilitating transfection of viral DNA [40] and has also been recently used to condense siRNA to nanoparticles, providing more efficient transfection [45]. Recently, calcium has been used to facilitate CPP-NA nanoparticle transfection, and it was shown that adding calcium provided highly efficient delivery of low toxicity [41-43].

Based on earlier research, we decided to assess the ability of calcium to facilitate CPP-based transfection of splice-correcting oligonucleotide (SCO) and siRNA. As a vector, we mostly used PepFect14 (PF14) – a CPP that was introduced in 2011 [26] and was shown to efficiently deliver siRNA [69], miRNA [7], SCO [26] and pDNA [70]. We assessed ion-complemented particles in three different cell lines: HeLa pLuc 705, HeLa EGFP 654 and U87-luc2. In addition to calcium, we tested other divalent metal ions found in living organisms, from which magnesium showed the highest efficiency, being tolerated by cells just as well as calcium, unlike some other divalent metal ions, e.g. zinc and manganese.

Nanoparticles of 100 nM SCO-705 and 500 nM PF14 had a hydrodynamic diameter of about 231 nm and a zeta potential of +9.3 mV. Adding calcium and magnesium ions at final concentrations of 0.3 or 3 mM did not significantly influence the size and charge of the nanoparticles.

Application of calcium-complemented PF14-SCO nanoparticles in Kole’s assay led to a strong increase in luminescence from HeLa pLuc 705 reporter cells. The ion-complemented nanoparticles provided a biological effect of SCO that is up to 10 times higher than without the ions. Based on luminescence change, we chose 0.3 mM and 3 mM as representative concentrations of the ions and used them in the following experiments. PCR analysis of the luciferase gene showed a substantial increase in the amount of the correctly spliced form of mRNA in the cells upon complementing SCO-PF14 nanoparticles with calcium and magnesium ions. These results confirm that addition of calcium and magnesium increases efficiency of splicing correction by the nanoparticles, which, in turn, translates to higher levels of the protein.

Interestingly, flow cytometry showed no significant increase in transfection efficiency of fluorescently labeled SCO into HeLa pLuc 705 cells upon addition of calcium and magnesium to the nanoparticles. However, there was a strong increase of the expression of EGFP as a result of splicing correction in HeLa EGFP 654 cells. It is in line with microscopy results that showed only a slight increase in transfection efficiency, but revealed remarkably higher nuclear localization of SCO-Cy5 that was transfected with calcium-complemented nanoparticles, and much stronger EGFP signal in cells transfected by ion-complemented PF14-SCO-654 nanoparticles. That gives reason to think that calcium and magnesium act mainly via facilitating endosomal escape of PF14-SCO nanoparticles, providing higher nuclear localization of oligonucleotide and, consequently, more efficient target-pre-mRNA binding, that results in more efficient splicing correction by the SCO.

We also investigated the role of class A scavenger receptors (SR-As) in transfection of ion-complemented nanoparticles. SR-As have been previously shown to play a major role in CPP-mediated transfection [71] and specifically in the uptake of PF14-containing nanoparticles [28, 36] by HeLa cells. We showed that these receptors are important also for the uptake of PF14-SCO-Ca^2+^ nanoparticles, as their blockage led to significant decrease of splicing correction efficiency by SCO. However, higher concentrations of SR-A inhibitors are required when calcium is added to the nanoparticles. It indicates a change in surface binding of ion-complemented nanoparticles: such particles can bind the receptors more strongly, competing more successfully with the inhibitors than non-complemented nanoparticles, or can be internalized via other routes.

Considering that calcium and magnesium act most likely via facilitating endosomal escape of SCO-PF14 nanoparticles, we pre-incubated HeLa pLuc 705 cells with endosome-destabilizing compound chloroquine in order to see if ion-complemented and non-complemented nanoparticles are equally affected by it. Our results showed that while chloroquine provided a 9-fold increase in efficiency of SCO-PF14 nanoparticles, this difference was lower upon addition of calcium and magnesium. The effect was ion concentration-dependent, and chloroquine provided only 1.5-fold increase of luciferase expression when 3 mM calcium was added. It confirms that the main feature of ion-complemented nanoparticles that explains their enhanced efficiency is most probably their greater ability to facilitate endosomal escape.

In addition to PF14, we tested several other peptides from PepFect and NickFect families. Our results showed that the effect observed in the case of PepFect14 is not necessarily the same for all cell-penetrating peptides. While providing a significant increase of biological activity of SCO transfected by some peptides – e.g., PF14 [26], C22-PF14 [22] and NF55 [65] – addition of calcium did not have any beneficial effect on transfection by others, e.g., NF70 [39], NF72 [39] and PF6 [25]. Remarkably, all three latter CPPs were designed as pH-sensitive delivery vehicles for siRNA. PF6 has four chloroquine-like residues in its structure, and NF70 and NF72 contain multiple histidine residues in their sequence to provide proton-sponge effect and facilitate escape from endosomes. This further corroborates that calcium ions mostly facilitate endosomal escape and cannot increase the efficacy of CPPs swiftly breaking out from endosomes.

It was previously shown that calcium can condense siRNA to nanoparticles [45] and facilitate its delivery with CPPs [42], providing more efficient endosomal escape and stronger gene silencing. We also tested calcium and magnesium in a siRNA-based gene silencing system, namely in the luciferase-expressing U87-luc2 cell line. Calcium significantly improved efficiency of siRNA-PF14 nanoparticles in the case of all siRNA concentrations applied, as well as of siRNA alone, in concordance with earlier research. Upon addition of magnesium, a significant increase of efficiency was observed only for siRNA-PF14 nanoparticles, but not for siRNA alone. It gives reason to suggest that, in the case of siRNA, mechanisms of action of calcium and magnesium are considerably different. It can translate into different efficiency of cargo, as in the case of SCO-PF14-Ca^2+^/Mg^2+^ nanoparticles, or into complete absence of the effect in the case of one of the ions, as in the case of siRNA-Mg^2+^ nanoparticles.

We presumed that enclosure of calcium and magnesium ions at higher concentrations could reduce the viability of cells treated with such nanoparticles. However, no impairment of cell viability was observed neither with high – up to 10 mM – concentrations of calcium and magnesium, nor in the case of incubation with nanoparticles containing up to 2 µM PF14. It indicates that nanoparticles that are used in our experiments – containing 500 nM peptide and up to 3 mM calcium or magnesium ions – are completely safe for cells.

Altogether, our results show that adding calcium and magnesium ions is an easy and non-toxic way to improve efficiency of different types of CPP-mediated transfection. It can be a promising approach as calcium and magnesium are present in the cells and in an extracellular environment at relatively high concentrations, being „natural” for cells and not causing an immune response. Resulting nanoparticles are easy-to-prepare, have physiologically safe diameter and lack cytotoxicity, at the same time providing several-fold increase of nanoparticle efficiency.

## Acknowledgements

We thank Piret Arukuusk for NickFect, and Tõnis Lehto for PF14-C22 peptides, and Kaida Koppel for preparation of TEM specimens. We highly appreciate help of Dmitri Lubenets at performing CLSM and FACS experiments.

This work was supported by the Estonian Ministry of Education and Research (0180019s11), Estonian Research Council (PUT1617P, PRG1169, PRG1259), Institute of Technology basic financing grant (PLTTI20912) to MP, and Leo Foundation (LF17040 to AR).

## Author Contributions

Maria Maloverjan: investigation, validation, visualization, writing – original draft. Kärt Padari: methodology, validation, visualization. Aare Abroi: investigation, validation. Ana Rebane: conceptualization, validation. Margus Pooga: conceptualization, methodology, funding acquisition, project administration, supervision, validation, writing – review & editing.

## Conflicting interests

Authors declare no conflict of interest.

---

**STable 1.**
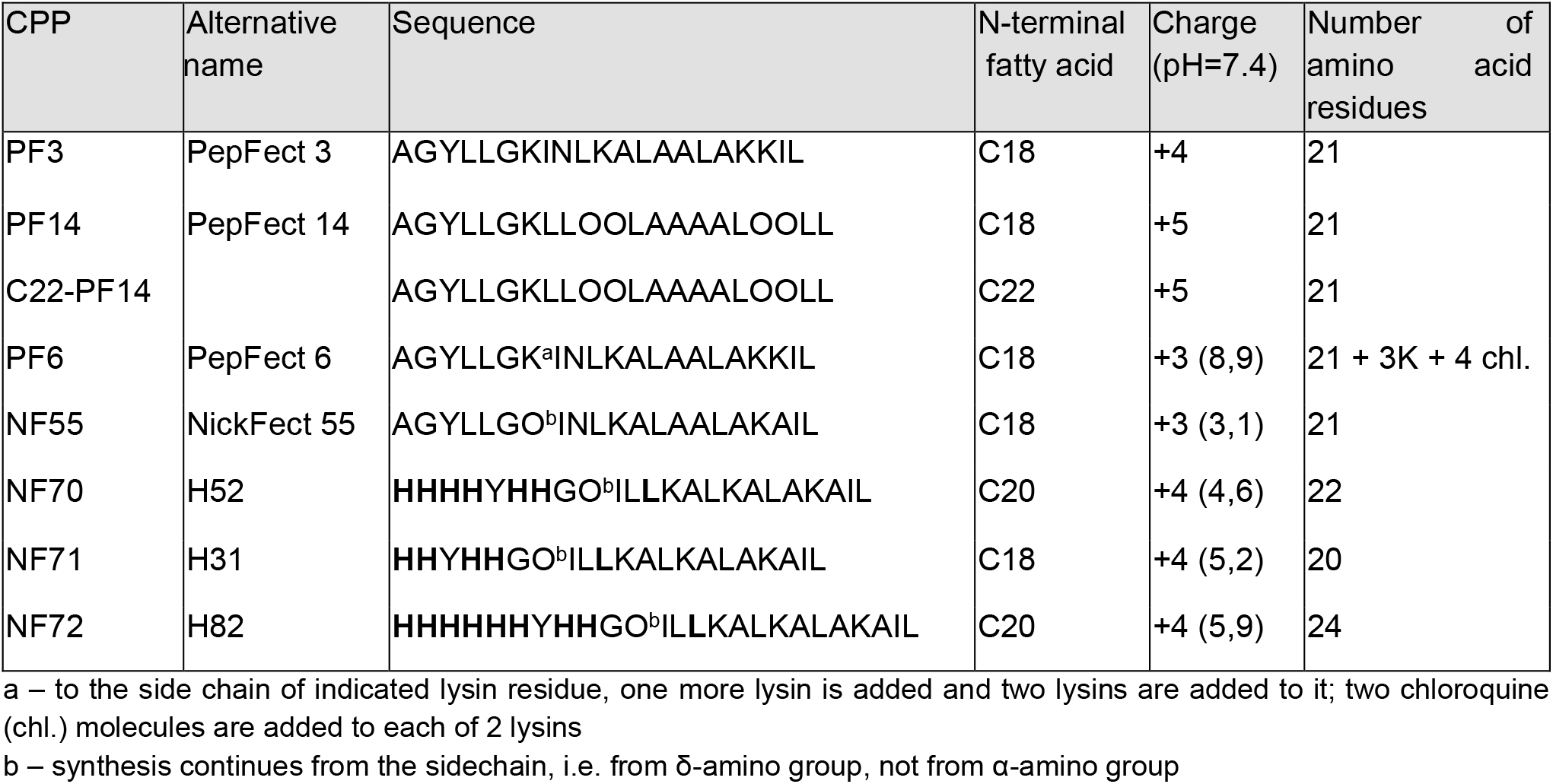
CPPs used, their sequences, charges and number of amino acid residues. All properties of the peptides are modelled and charges or logD values are calculated using MarvinSketch 15.9.14, ChemAxon Ltd, USA (see [39]).

**SFig. 1.**
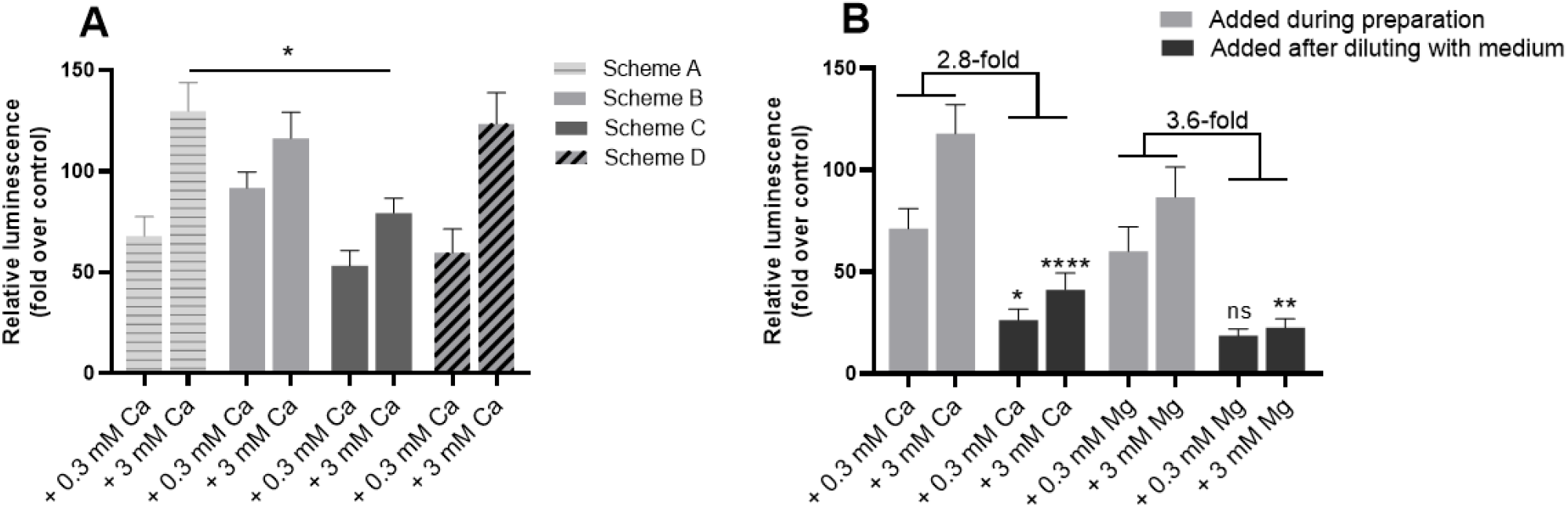
Effect of SCO-PF14 nanoparticles on luciferase expression in case of different preparation schemes. HeLa pLuc 705 cells were incubated with the solutions containing differently prepared nanoparticles of 100 nM SCO, 500 nM PF14 and 0.3 or 3 mM Ca^2+^. Scheme A: mixing SCO with PF14, adding Ca^2+^ after 15 min. Scheme B: mixing SCO with PF14 and Ca^2+^. Scheme C: mixing PF14 with Ca^2+^, adding SCO after 15 min. Scheme D: mixing SCO with Ca^2+^, adding PF14 after 15 min. For all schemes, overall incubation time before diluting the nanoparticle-containing solutions with cell culture medium was 30 min. Luminescence was measured after 24 h with microplate reader Infinite M200 PRO (Tecan, Switzerland). Each dataset represents mean + SEM of 4 independent experiments. Data was analyzed by one-way ANOVA with Tukey’s test, *p-value < 0.05, **p-value < 0.005, ****p-value < 0.0001, ns – not significant.

**SFig. 2.**
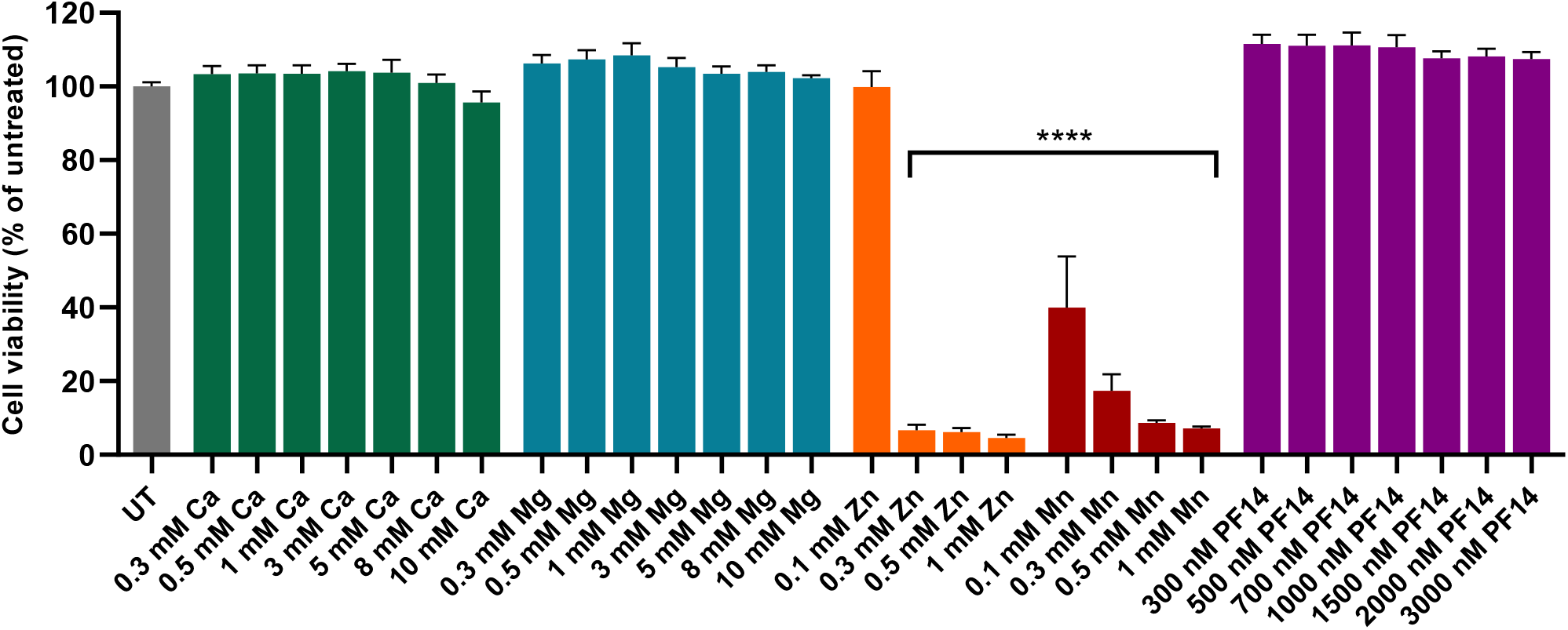
The effect of PF14 and divalent metal ions on viability of cells. HeLa pLuc 705 cells were incubated for 24 h with solutions containing PF14, Ca^2+^, Mg^2+^, Zn^2+^ or Mn^2+^ at different concentrations. Viability of the cells was evaluated using WST-1 assay, and absorption of untreated cells (UT) was taken for 100%. Each dataset represents mean + SEM of at least 3 independent experiments. Data was analyzed by one-way ANOVA with Dunnett’s test, asterisks indicate statistically significant difference compared to untreated cells, ****p-value < 0.0001.

## Notes

### Competing Interest Statement

The authors have declared no competing interest.

